# Including phenotypic causal networks in genome-wide association studies using mixed effects structural equation models

**DOI:** 10.1101/251421

**Authors:** Mehdi Momen, Ahmad Ayatollahi Mehrgardi, Mahmoud Amiri Roudbar, Andreas Kranis, Renan Mercuri Pinto, Bruno D. Valente, Gota Morota, Guilherme J. M. Rosa, Daniel Gianola

## Abstract

**Background:** Phenotypic networks describing putative causal relationships among multiple phenotypes can be used to infer single-nucleotide polymorphism (SNP) effects in genome-wide association studies (GWAS). In GWAS with multiple phenotypes, reconstructing underlying causal structures among traits and SNPs using a single statistical framework is essential for understanding the entirety of genotype-phenotype maps. A structural equation model (SEM) can be used for such purposes.

**Methods:** We applied SEM to GWAS (SEM-GWAS) in chickens, taking into account putative causal relationships among body weight (BW), breast meat (BM), hen-house production (HHP), and SNPs. We assessed the performance of SEM-GWAS by comparing the model results with those obtained from traditional multi-trait association analyses (MTM-GWAS).

**Results:** Three different putative causal path diagrams were inferred from highest posterior density (HPD) intervals of 0.75, 0.85, and 0.95 using the inductive causation algorithm. A positive path coefficient was estimated for BM→BW, and negative values were obtained for BM→HHP and BW→HHP in all implemented scenarios. Further, the application of SEM-GWAS enabled the decomposition of SNP effects into direct, indirect, and total effects, identifying whether a SNP effect is acting directly or indirectly on a given trait. In contrast, MTM-GWAS only captured overall genetic effects on traits, which is equivalent to combining the direct and indirect SNP effects from SEMGWAS.

**Conclusions:** Although MTM-GWAS and SEM-GWAS use the same probabilistic models, we provide evidence that SEM-GWAS captures complex relationships and delivers a more comprehensive understanding of SNP effects compared to MTM-GWAS. Our results showed that SEM-GWAS provides important insight regarding the mechanism by which identified SNPs control traits by partitioning them into direct, indirect, and total SNP effects.

## Background

Genome-wide association studies (GWAS) have become a standard approach for investigating relationships between common genetic variants in the genome (e.g., single-nucleotide polymorphisms, SNPs) and phenotypes of interest in human, plant, and animal genetics [1–4]. A typical GWAS is based on univariate linear or logistic regression of phenotypes on genotypes for each SNP individually while often adjusting for the presence of nuisance covariates [5]. A statistically significant association indicates that SNPs may be in strong linkage disequilibrium (LD) with quantitative trait loci (QTLs) that contribute to the trait etiology. Alternatively, multi-trait model GWAS (MTM-GWAS) can be used to test for genetic associations among a set of traits [6–8]. It has been established that MTM-GWAS reduces false positives and increases the statistical power of association tests, explaining the recent popularity of this method. MTM-GWAS can be used to study genetic associations of multiple traits; however, it does not identify factors that mediate relationships between the detected effects and dependencies involving complex traits.

Complex traits are the product of various cryptic biological signals that may affect a trait of interest either directly or indirectly through other intermediate traits [9]. A standard regression cannot describe such complex relationships between traits and QTLs properly. For instance, some traits may simultaneously act as both dependent and independent variables. Structural equation modeling (SEM) is an extended version of Wright’s path analysis [10, 11] that offers a powerful technique for modeling causal networks. In a complex genotype-phenotype setting involving many traits, a given trait can be influenced not only by genetic and systematic factors but also by other traits (as covariates) as well. Here, QTLs may not affect the target trait directly; instead, the effects may be mediated by upstream traits in a causal network. Indirect effects may therefore constitute a proportion of perceived pleiotropy, and these concepts apply to sets of heritable traits, organized as networks, that are common in biological systems. An example from dairy cattle production systems, described by Gianola and Sorensen [11], is that higher milk yield increases the risk of a particular disease, such as mastitis, while the prevalence of the disease may negatively affect milk yield As another example, Varona, et al. [12] explored a causal link from litter size to average piglet weight in two pig breeds. In humans, obesity is a key factor influencing insulin resistance, which subsequently causes type 2 diabetes. Lists of causal networks across human diseases and candidate genes are described in Kumar and Agrawal [13] and Schadt [14].

Although MTM-GWAS is a valuable approach, it only captures correlations or associations among traits and does not provide information about causal relationships. Knowledge of the causal structures underlying complex traits is essential, as correlation does not imply causation. For example, a correlation between two traits, T1 and T2, could be attributed to a direct effect of T1 on T2 or T2 on T1, or to additional variables that jointly influence both traits [15]. Likewise, if we know a “causal” SNP is linked to a QTL, we can imagine three possible scenarios: 1) causal (*SNP* → *T*1 → *T*2), 2) reactive (*SNP* → *T*2 → *T*1), or 3) independent (*T*1 ← *SNP* → *T*2). Scenarios (1) and (2) do not cause pleiotropy but produce association.

A SEM methodology has the ability to handle complex genotype-phenotype maps in GWAS, placing an emphasis on causal networks [16]. Therefore, SEM-based GWAS (SEM-GWAS) may provide a better understanding of biological mechanisms and of relationships among a set of traits than MTM-GWAS. SEM can potentially decompose the total SNP effect on a trait into direct and indirect (i.e., mediated) contributions. However, SEM-derived GWAS has yet not been discussed or applied fully in quantitative genetic studies yet. Our objective was to illustrate the potential utility of SEM-GWAS by using three production traits in broiler chickens genotyped for a battery of SNPs as a case example.

## Methods

### Data set

The analysis included records for 1,351 broiler chickens provided by Aviagen Ltd. (Newbridge, Scotland) for three phenotypic traits: body weight (BW), ultrasound of breast muscle (BM) at 35 days of age, and hen-house egg production (HHP), defined as the total number of eggs laid between weeks 28 and 54 per bird. The sample consisted of 274 full-sib families, 326 sires, and 592 dams. More details regarding population and family structure were provided by Momen, et al. [17]. A pre-correction procedure was performed on the phenotypes to account for systematic effects such as sex, hatch week, pen, and contemporary group for BW, BM, and HHP.

Each bird was genotyped for 580,954 SNP markers with a 600k Affymetrix SNP [18] chip (Affymetrix, Inc., Santa Clara, CA, USA). The Beagle software program [19] was used to impute missing SNP genotypes, and quality control was performed using PLINK version 1.9 [20]. After removing markers that did not fulfill the criteria of minor allele frequencies < 1%, call rate > 95%, and Hardy–Weinberg equilibrium (Chi-square test p-value threshold was 10^− 6^), 354,364 autosomal SNP markers were included in the analysis.

### Multiple-trait model for GWAS

MTM-GWAS is a single-trait GWAS model extended to multi-dimensional responses. When only considering additive effects of SNPs, the phenotype of a quantitative trait using the single-trait model can be described as:

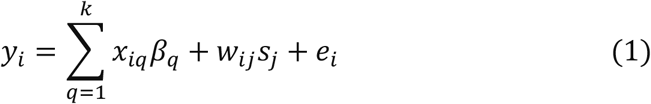

where *y_i_* is the phenotypic trait of individual *i*, *x_iq_* is the incidence value for the *i*th phenotype in the *q*th level of systematic environmental effects, *β_q_* is the fixed effect of the *q*th systemic environmental effect on the trait, *w_j_* = (*w*_1_, …, *w_p_*) is the number of A alleles (i.e., *w_j_* ∈ {0,1, 2}) in the genotype of SNP marker *j*, and *s*_j_ is the allele substitution effect for SNP marker *j*. Strong LD between markers and QTLs coupled with an adequate marker density increases the chance of detecting marker and phenotype associations. Hypothesis testing is typically used to evaluate the strength of the evidence of a putative association. Typically, a *t*-test is applied to obtain p-values, and the statistic is *T*_ij_ = *ŝ*_j_/*se*(*ŝ*_j_), where *ŝ* is the point estimate of the *j*th SNP effect and *se*(*ŝ*_j_) is its standard error.

The single locus model described above is naïve for a complex trait because the data typically contain hidden population structure and individuals have varying degrees of genetic similarity [21,22]. Therefore, accounting for covariance structure induced by genetic similarity is expected to produce better inferences [23]. Ignoring effects that reveal genetic relatedness inflates the residual terms and compromises the ability to detect association. A random effect *g*_*i*_, including a covariance matrix reflecting pairwise similarities between additive genetic effects of individuals, can be included to control population stratification. The similarity metrics can be derived from pedigree information or from whole-genome marker genotypes. This model, extended for analysis of *t* traits, is given by:

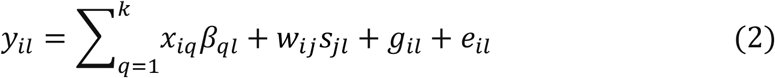

for *i* = 1,2,···, *n,l* = 1,2,···, *t*. In this extension, *y_il_* is the phenotypic value of the *l*th trait for the *i*th subject, *β_qi_* is the systematic effect of the *q*th environmental factor *x_iq_* on the *l*th trait, *s_jl_* is the additive effect of the *j*th marker on the *l*th trait, *w_ij_* is as previously defined, and *g_il_* and *e_il_* are the random polygenic effect and model residual assigned to individual *i* for trait *l*, respectively. Random effects within a trait follow the multivariate normal distribution, 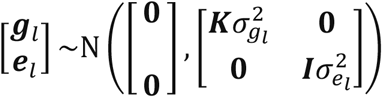, where ***K*** is a genetic relationship matrix, 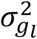 is the additive genetic variance of trait *l, **I*** is an identity matrix, and 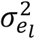 is the residual variance for trait *l*. The multiple-trait model accounts for the additive genetic (*ρ_ll’_*) and residual correlation (*λ_ll′_*) between a pair of traits *l* and *l*’.

The positive definite matrix **K** may be a genomic relationship matrix (**G**) computed from marker data, or a pedigree-based matrix (**A**) computed from genealogical information. The **A** matrix describes the expected additive similarity among individuals, while **G** measures the realized fraction of alleles shared. Genomic relationship matrices can be derived in several ways [24–26]. Here, we used the form proposed by VanRaden 2008 [24]:

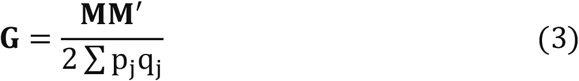

where **M** is an *n × p* matrix of centered SNP genotypes and *p_j_* and *q_j_* = 1 − *p_j_* are the allele frequencies at marker locus *j*. We evaluated both **A** and **G** in the present study.

### Structural equation model association analysis

A SEM consists of two essential parts: a measurement model and a structural model. The measurement model depicts the connections between observable variables and their corresponding latent variables. The measurement model is also known as confirmatory factor analysis. The critical part of a SEM is the structural model, which can have three forms. The first consists of observable exogenous and endogenous variables. This model is a restricted version of a SEM known as path analysis [10]. The second form explains the relationship between exogenous and endogenous variables that are only latent. The third type is a model consisting of both manifest and latent variables.

SEM can be applied to GWAS as an alternative to MTM-GWAS to study how different causal paths mediate SNP effects on each trait. The following SEM model was considered:

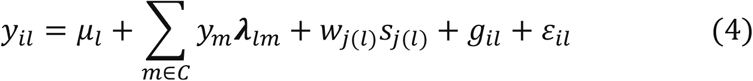

where *C* is the set of phenotypic traits that directly affect the trait *l, ***λ***_lm_*is a structural coefficient representing the effect of trait *m* on trait *l*, and *g_l_*~*N*(0, ***K***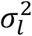) is the polygenic effect of the *l*th trait. The remaining terms are as presented earlier with one important difference: the SNP effects are not interpreted as overall effects on trait *l* but instead represent direct effects on trait *l*. Additional indirect effects from the same SNP may be mediated by phenotypic traits in *C*. Each marker is entered into equation (4) one at a time, and its significance is tested. For a discussion of how SEM represents genetic signals on each trait through multiple causal paths, see Wu et al. [27] and Jamrozik and Schaeffer [28]. Despite the difference in interpretation, the distribution of the vector of polygenic effects is assumed to be the same as in the MTM-GWAS model. The same applies to residual terms within a trait. We also consider trait-specific residuals to be independent within an individual. This restriction is required to render structural coefficients likelihood-identifiable. In addition, the interpretation of inferences as having a causal meaning requires imposing the restriction that the residuals’ joint distribution be interpreted as the causal sufficiency assumption [29]. In the present study, all exogenous and endogenous variables were observable, and there was no latent variable. Hence, causal structure was assumed between the endogenous variables BM, BW, and HHP.

We considered the following GWAS models, which their causal structures were recovered by the inductive causation (IC) algorithm [29]: (1) MTM-GWAS with pedigree-based kinship **A** (MTMA) or marker-based kinship **G** (MTM-G), and (2) SEM-GWAS with **A** (SEM-A) or **G** (SEM-G). Although nuisance covariates such as environmental factors can be omitted in the graph, they may be incorporated into the models as exogenous variables. The SEM representation allowed us to decompose SNP effects into direct, indirect, and total effects.

A direct SNP effect is the path coefficient between a SNP as an exogenous variable and a dependent variable without any causal mediation by any other variable. The indirect effects of a SNP are those mediated by at least one other intervening endogenous variable. Indirect effects are calculated by multiplying path coefficients for each path linking the SNP to an associated variable, and then summing over all such paths [30]. The overall effect is the sum of all direct and indirect effects. By explicitly accounting for complex relationship structure among traits in such a way, SEM provides a better understanding of a genome-wide SNP analysis by allowing us to decompose effects into direct, indirect, and overall effects within a predefined casual framework [31]. MTMGWAS and SEM-GWAS were compared with the logarithm of the likelihood function (log L), Akaike’s Information Criterion (AIC), and the Bayesian Information Criterion (BIC). The model providing the lowest values for these information criteria is considered to fit the data better [27]. MTM-GWAS and SEM-GWAS were fitted using the SNP Snappy strategy, which is implemented in the Wombat software program [32].

### Searching for a phenotypic causal network in a mixed model

In the SEM-GWAS formulation described earlier, the structure of the underlying causal phenotypic network needs to be known. Because this is not so in practice, we used a causal inference algorithm to infer the structure. Residuals are assumed to be independent in all SEM analyses, so associations between observed traits are viewed as due to causal links between traits and by correlations among genetic values (i.e., *g*_1_, *g*_2_, and *g*_3_). Thus, to eliminate confounding problems when inferring the underlying network among traits, we used the approach of Valente, et al. [32] to search for acyclic causal structures through conditional independencies on the distribution of the phenotypes, given the genetic effects. A causal phenotypic network was inferred in two stages: 1) an MTM model [33] was employed to estimate covariance matrices of additive genetic effects and of residuals, and 2) the causal structure among phenotypes from the covariance matrix between traits, conditionally on additive genetic effects, was inferred by the IC algorithm. The residual (co)variance matrix was inferred using Bayesian Markov-chain Monte Carlo [27, 32], with samples drawn from the posterior distribution. The reason for our use of the residual (co)covariances is that the residual structure could bear information from the joint distribution of all phenotypic traits conditional on their polygenic effects, such that they correct the confounding issues caused by such effects when the traits are genetically correlated [29]. For each query testing statistical independence between traits *y_l_* and *y_l′_*, the posterior distribution of the residual partial correlation *ρ_yl,y_1_′_* |*S* was obtained, where *S* is a set of variables (traits) that are independent. Three highest posterior density (HPD) intervals of 0.75, 0.85, and 0.95 were used to make statistical decisions for SEM-GWAS. We thus considered SEM-A75 (HPD > 0.75), SEM-A85 (HPD > 0.85), SEM-A95 (HPD > 0.95), and SEMG75 (HPD > 0.75). An HPD interval that does not contain zero declares *y_l_* and *y_l′_* to be conditionally dependent.

## Results

Figure 1 shows phenotypic relationship structures recovered by the IC algorithm for the three different HPD intervals. Edges connecting two traits represent non-null partial correlations as indicated by HPD intervals. We compared the two MTM-GWAS and four SEM-GWAS by using the three chicken traits (BW, BM, and HHP). Only causal structures among the three traits are shown in Figure 1, because other parts were the same across the different SEM models. Fully recursive SEM-A75 and SEM-G75 revealed direct effects of BM on BW and HHP, and those of BW on HHP, as well as an indirect effect of BM on HHP. In addition, SEM-A85 detected a direct effect of BM on BW, the direct effect of BW on HHP, and the indirect effect of BM on HHP mediated by BW. Finally, SEM-A95 only identified a direct effect of BM on BW because of a statistically stringent HPD cutoff imposed.

Given the causal structures inferred from the IC algorithm, the following SEM was fitted:

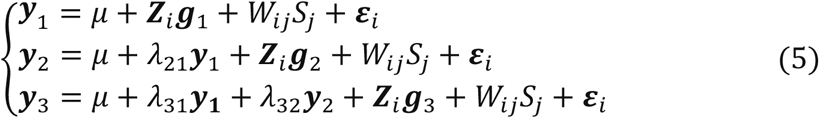

Note that only a small number of the entries in the structural coefficient matrix (*λ* in equation 5) are nonzero due to sparsity. These nonzero entries specify the effect of one phenotype on other phenotypes. The corresponding directed acyclic graph is shown in Figure 2 assuming the causal relationships among the three traits, where y_1_, *y*_2_, and *y*_3_ represent BM, BW, and HHP, respectively; *SNP*_j_ is the genotype of the *j*th SNP; *S_jl_* is the direct SNP effect on trait *l*; and the remaining variables are as presented earlier. This diagram depicts a fully recursive structure in which all recursive relationships among the three phenotypic traits are shown. Arrows represent causal connections, whereas double-headed arrows between polygenic effects are correlations.

**Figure 1.**
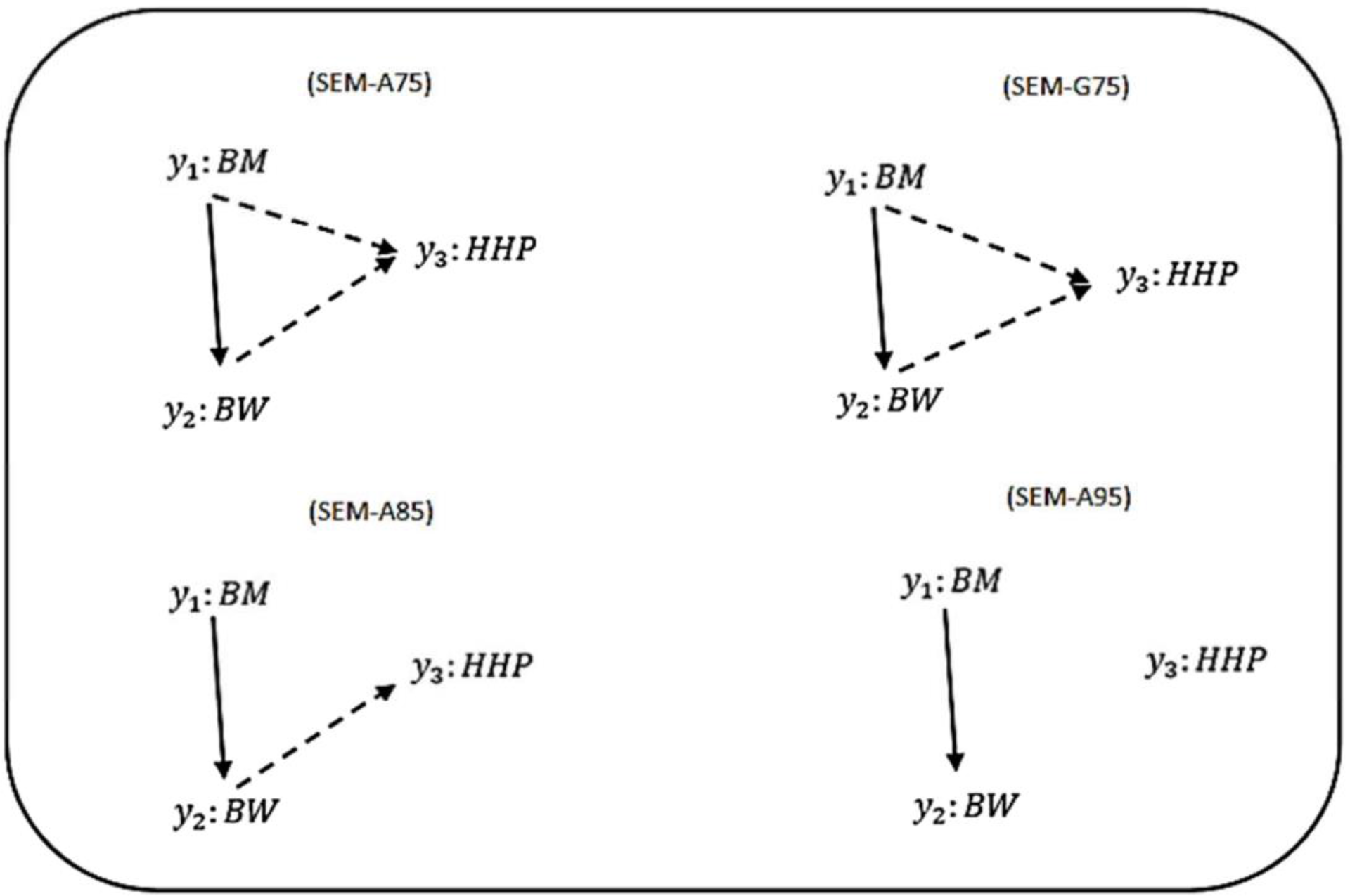
Causal graphs inferred using the IC algorithm among three traits: breast meat (BM), body weight (BW) and hen-house production (HHP) in the chicken data. SEM-A75 and SEM-G75 were the inferred fully recursive causal structures with HPD > 0.75 and corrected for genetic confounder using **A** (pedigree-based) and **G** (marker-based) matrices. SEM-A85 and SEMA95 were obtained with HPD > 0.85 and HPD > 0.95, respectively, corrected with **A**. Arrows indicate direction of causal relationships. Dashed lines indicate negative coefficients, and the continuous arrows indicate positive coefficients.

**Figure 2.**
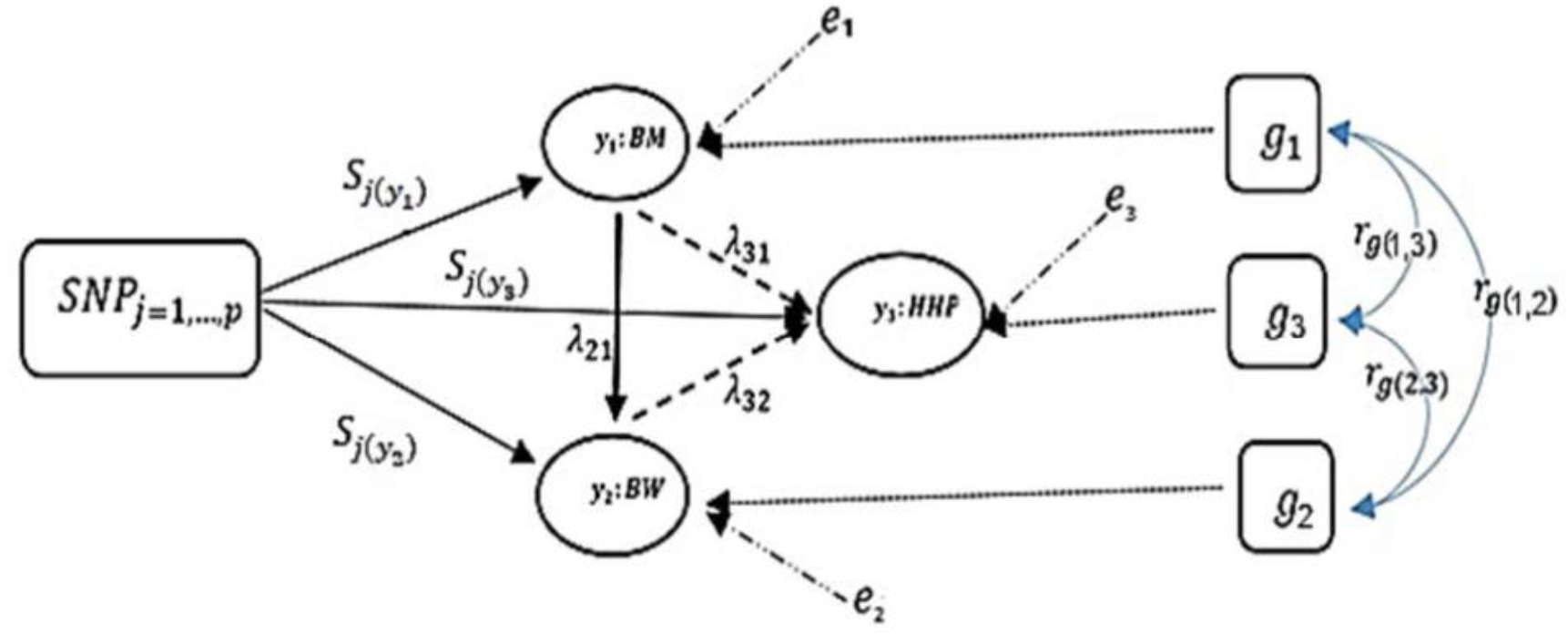
A diagram for causal path analysis of SNP effects in a fully recursive structural equation model for three traits, *p* exogenous independent SNP variables, and three correlated polygenic effects. Arrows indicate the direction of causal effects and dashed lines represent associations among the three phenotypes. Genetic correlation between traits (*r_g_*), polygenic effects (*g_l_*), environmental effect on trait *l* (*e_l_*), effects of *j* th SNP on *l* th trait (*S_j(yl)_*), and recursive effect of phenotype *l*′ on phenotype *l* (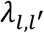).

We examined the fit of each model implemented to assess how well it describes the data (Table 1). Varona, et al. [12] and recently Valente, et al. [34] showed that re-parametrization and reduction of a SEM mixed model yield the same joint probability distribution of observation as in MTM, suggesting that the expected likelihood of SEM and MTM should be the same. As expected, SEMGWAS and MTM-GWAS showed very similar results (e.g., SEM-A75 vs. MTM-A and SEM-G75 vs. MTM-G). Among the models considered, those involving **G** exhibited slightly better fits. SEMA85 and SEM-A95, sharing a subset of the SEM-A75 structure, presented almost identical AIC and BIC values. Since these results imply that the recursive model and standard mixed model for GWAS are statistically equivalent in terms of the fitting criteria, the focus of the remainder of the analysis will be on the modeling of SNP (or QTL) effects in the SEM context as an extension of MTM, which accounts for recursive links among the three measured traits.

**Table 1.**
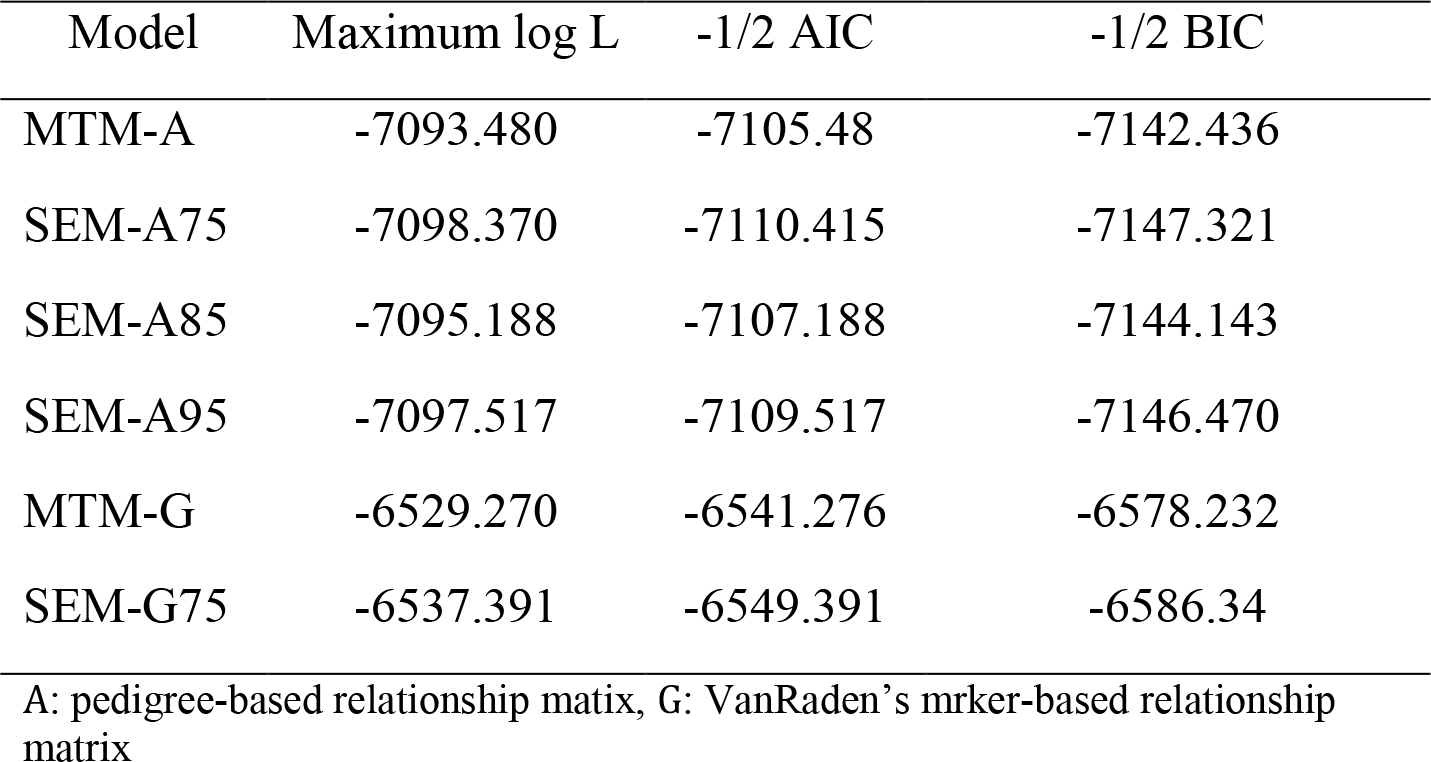
Model comparison criteria: logarithm of the restricted maximum likelihood function (log L), Akaike’s information criteria (AIC), Schwarz Bayesian information criteria (BIC) to evaluate model fit for two MTM and four SEM models.

### Structural coefficients

Table 2 presents the causal structural path coefficients for endogenous variables (BM, BW, and HHP). All models have positive effects for BM→BW, whereas the BM→HHP and BW→HHP relationships have negative path coefficients. The latter confirmed the fact that chicken breeding is divided into broiler and layer sections due to the negative genetic correlation between BW and HHP.

**Table 2.** Estimates of three causal structural coefficients (*λ*) derived from four different structural models. BM: breast meat. BW: body weight. HHP: hen-house production. SEM-75: HPD > 0.75. SEM-G75: HPD > 0.75. SEM-A85: HPD > 0.85. SEM-A95: HPD > 0.95.

| Path | Structural Models |  |  |  |
| --- | --- | --- | --- | --- |
|  | SEM-A75 | SEM-G75 | SEM-A85 | SEM-A95 |
| $\lambda_{BM \rightarrow BW}(\lambda_{21})$ | 2.13 | 2.19 | 2.14 | 2.14 |
| $\lambda_{BM \rightarrow HHP}(\lambda_{31})$ | -0.17 | -0.28 | *** | *** |
| $\lambda_{BW \rightarrow HHP}(\lambda_{32})$ | -0.27 | -0.096 | -0.31 | *** |

Also shown in Table 2 are the magnitudes of the SEM structural coefficient reflecting the intensity of the causality. The positive coefficient *λ*_21_ quantifies the (direct) causal effect of BM on BW. This suggests that a 1-unit increase in BM results in a *λ*_21_-unit increase in BW. Likewise, the negative causal effects *λ*_31_ and *λ*_32_ offer the same interpretation.

### Decomposition of SNP effect paths using a fully recursive model

We can decompose SNP effects into direct and indirect effects using Figure 2. The direct effect of the SNP *j* on *y*_3_ (HHP) is given by 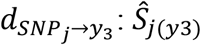, where *d* denotes the direct effect. Note there are only one direct and many indirect paths. We find three indirect paths from *SNP_j_* to *y*_3_ mediated by *y*_1_ and *y*_2_ (i.e., the nodes formed by other traits). The first indirect effect is 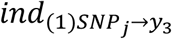: *λ*_32_(*λ*_21_*Ŝ_j_*_(*y*1)_) in the path mediated by y_1_ and y_2_, where *ind* denotes the indirect effect. The second indirect effect 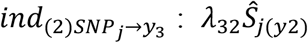, is mediated by *y_2_*. The last indirect effect, is 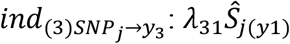, mediated by y_1_. Therefore, the overall effect is given by summing all four paths, 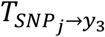: *λ*_32_(*λ*_21_*Ŝ_j_*_(*y*1)_) + *λ*_32_*Ŝ_j_*(*y*_2_) + *λ*_31_*Ŝ_j_*_(*y*1)_ + *Ŝ_j_*_(*y*3)_. The fully recursive model of the overall SNP effect is then:

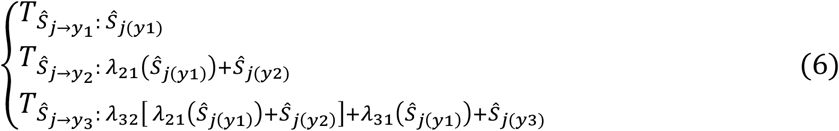

For *y*_1_ (BM), there is only one effect, so the overall effect is equal to the direct effect. For *y*_2_ (BW) and *y*_3_ (HHP), direct and indirect SNP effects are involved. There are two paths for *y*_2_: one indirect, 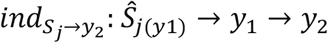, and one direct, *d_S_j_→y_2__:Ŝ_j(y2)_→y_2_*. Here, the SNP effect is direct and mediated thorough other phenotypes according to causal networks in SEM-GWAS (Figures 1 and 2). For instance, the overall SNP effect for *y*_3_ into four direct and indirect paths is 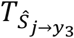 :*λ*_32_*λ*_21_*Ŝ_j_*_(*y*1)_ + *λ*_32_*Ŝ_j_*_(*y*1)_ + *λ*_31_*Ŝ_j_*_(*y*1)_ + *Ŝ_j_*_(*y*3)_.

The scatter plots in Figure 3 compare the estimated total effects for HHP (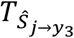) obtained from SEM-GWAS and those from MTM-GWAS. We observed good agreement between SEM-GWAS and MTM-GWAS. The total SNP signals derived from SEM and MTM are the same but SEM provides biologically relevant additional information.

**Figure 3.**
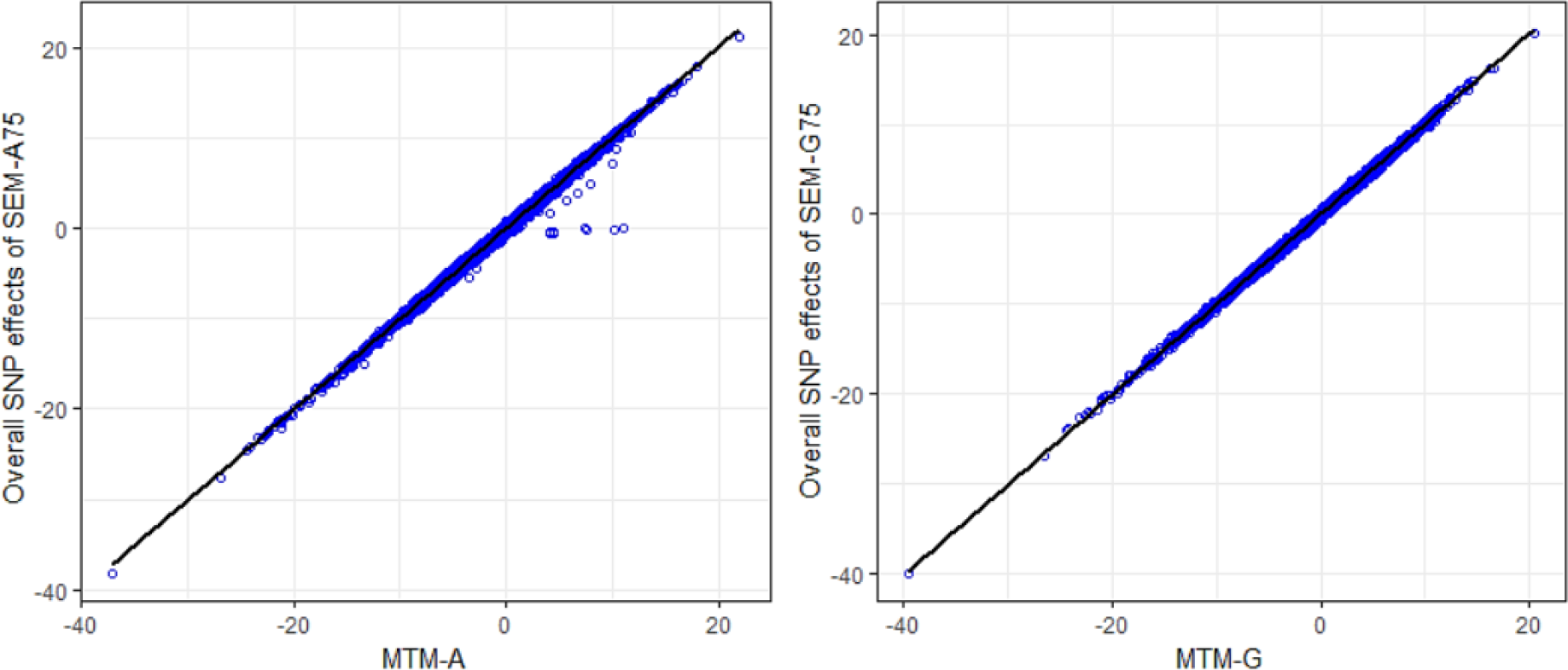
Comparison of multiple trait (MTM) and fully recursive overall SNP effects obtained with A (pedigree-based) and G (marker-based) from structural equation modeling (SEM)-based GWAS. Overall effects in SEM are the sum of all direct and indirect effects. HHP: hen-house egg production.

Supplementary Figures S1–S4 present scatter plots of MTM-GWAS and SEM-GWAS signals (SEM-A75, SEM-G75, SEM-A85, and SEM-A95) for the *BM → BW* path, which was a common path across all SEM-GWAS considered. These two traits have a genetic correlation of 0.5 (results not shown). We partitioned the SEM causal link into direct, indirect, and overall effects based on directed links inferred from the IC algorithm with HPD > 0.85, whereas MTM-GWAS captures an overall SNP effect on BW. Scatter plots of the overall effects from SEM-GWAS and those of the total effects from MTM-GWAS indicated almost perfect agreement (top left plots, Supplementary Figures S1–S4). We also observed concomitance between estimated overall and direct effects (top right plots, Supplementary Figures S1–S4). In contrast, there was less agreement in the magnitude of the SNP effects when comparing overall vs. indirect effects (bottom left plots, Supplementary Figures S1–S4). There was no linear relationship between the indirect and direct SNP effects (bottom right plots, Supplementary Figures S1–S4). In short, genetic signals detected in SEMGWAS were close to those of MTM-GWAS for overall effects because both models are based on a multivariate approach with the same covariance matrix. In all SEM-GWAS, results showed that direct effects contributed to overall effects more than the indirect effects.

### Manhattan plot of direct, indirect, and overall SNP effects

Figure 4 depicts a Manhattan plot summarizing the magnitude of direct (SEM-75A), indirect (SEM-75A), and overall SNP effects (MTM-75A). We plotted the decomposed SNP effects on BW along chromosomes to visualize estimated marker effects from SEM-GWAS and MTMGWAS. The indirect and direct effects provide a view of SNP effects from a perspective that is not available for the total effect of MTM-GWAS. For instance, many pleiotropic QTLs have positive direct effects on BW but negative effects on BM. There were two estimated SNP effects on chromosomes 1 and 2 that deserve particular attention. These two SNPs are highlighted with black circles and red ovals. The overall effect of the first SNP consisted of large indirect and small direct effects on BM, whereas the opposite pattern was observed for the second SNP, which showed large direct and small indirect effects. Although the overall effects of these SNPs were similar (top Manhattan plot, Figure 4), use of decomposition allowed us to determine that the trait of interest is affected in different manners: the second SNP effect acted directly on BW without any mediation by BM, whereas the first SNP reflected a large effect mediated by BM on BW. Collectively, new insight regarding the direction of SNP effects can be obtained using the SEM-GWAS methodology.

The corresponding Manhattan plot based on –log_10_ (p-values) is shown in Supplementary Figure S5. As with the magnitude of effect sizes, the results showed that –log_10_ (p-values) of estimated overall effects from SEM-A75 and those from MTM-A75 yielded the same significant peaks. We found that some significant indirect SNP effects reached genome-wide significance after correction for multiple-testing using a 5% FDR threshold level (2.752). The most significant SNPs were on chromosomes 1 and 4 (GGA1 and GGA4).

**Figure 4.**
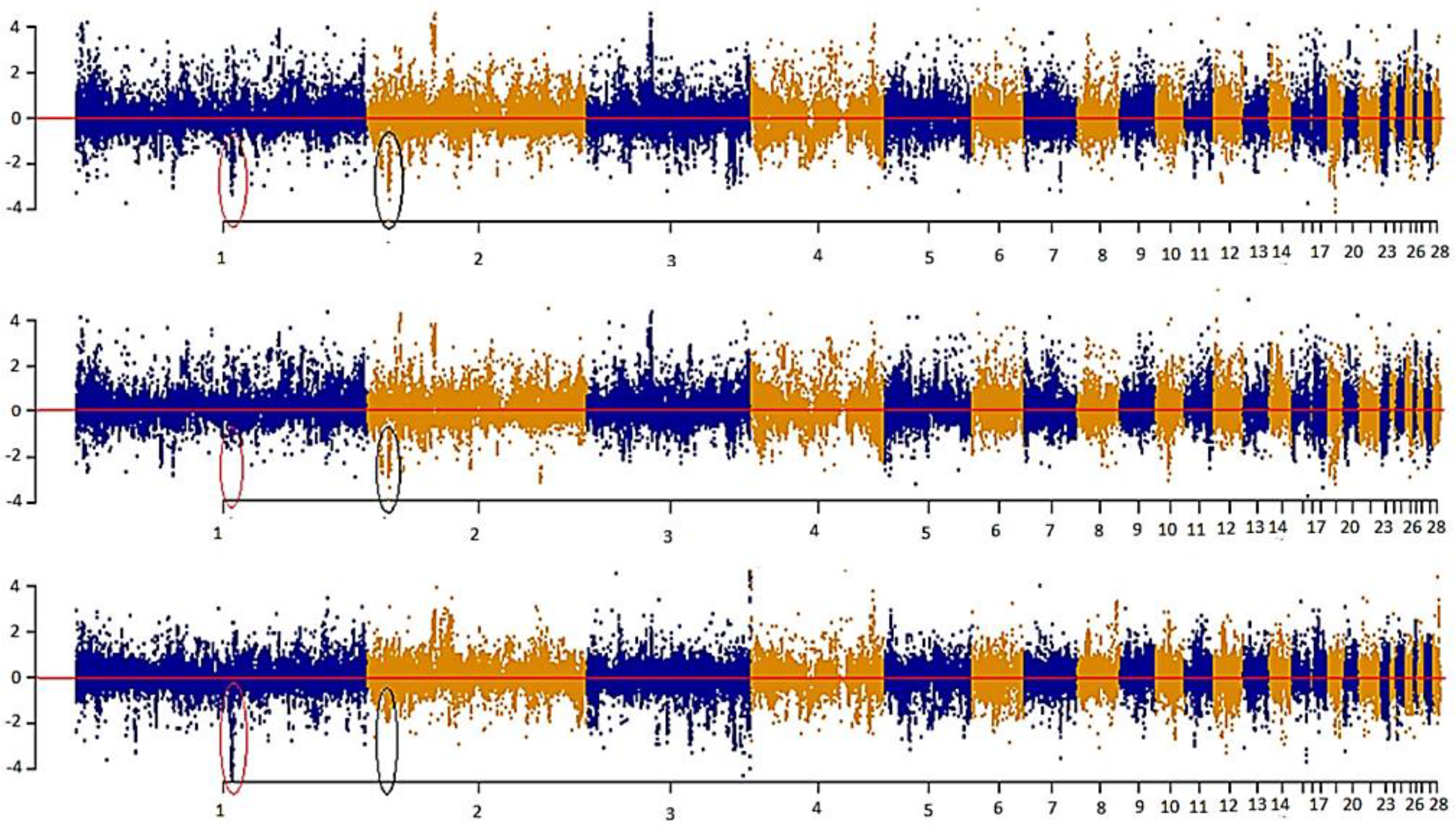
Manhattan plot showing overall, direct, and indirect SNP effects using a full recursive model based on A matrix for body weight (BW).

As an illustration, the six most significant SNPs with the highest –log_10_ (p-values) for each type of decomposed SNP effect are presented in Table 3. Seven candidate genes were identified near the significant SNPs derived from the SNP effects decomposition, with two on GGA7 (*OLA1* and *ZNF385B*), one on GGA3 (*EPHA7*), three on GGA4 (*LOC422264*, *LOC422265*, and *MAEA*), and one on GGA14 (*GRIN2A*). We found that only genes on GGA4 and GGA1 are linked to significant indirect SNP effects that impact HHP. Some studies reported QTLs for BM on GGA1 and for BW on GGA4, stating that these genomic regions contain QTLs related to abdominal fat and growth traits that were detected across diverse chicken populations [35, 36]. One of the two detected genes on GGA14, i.e., *GRIN2A*, which was linked to the SNP Gga_rs313620413, showed significant direct and overall SNPs effects using SEM as well as MTM. Collectively, Gga_rs15390496, Gga_rs16591372, and Gga_rs313620413 SNPs on GGA3, GGA7, and GGA14, which were linked to *EPHA7*, *OLA1*, and *GRIN2A*, respectively, represent candidate genes identified from overall effects of both SEM and MTM (Table 3).

We noted that the six SNPs selected according to the –log_10_ (p-values) from the direct effect on HHP (i.e., 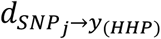) had small indirect effects ranging from –0.9018 to 0.2983. These indirect effects were negligible compared with their corresponding direct and total effects. Also, exploring the indirect effect sizes of the six most significant SNPs showed that indirect effects that are transmitted from inferred causal networks have the ability to change the magnitude of overall SNP effects, even changing them to the opposite direction (i.e., from positive to negative or vice versa).

It should also be noted that the estimated additive SNP effects obtained from the four SEM-GWAS can be used for inferring pleiotropy. For instance, a pleiotropic QTL may have a large positive direct effect on BW but may exhibit a negative indirect effect coming from BM, which in turn reduces the total QTL effect on BW. Arguably, the methodology employed here would be most effective when the direct and indirect effects of a QTL are in opposite directions. If the direct and indirect QTL effects are in the same direction, the power of SEM-GWAS may be the same as the overall power of MTM-GWAS. The overall effect (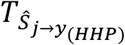) of a given SNP consisted of large indirect (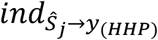) and small direct (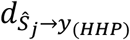) effects on HHP, as observed for the top most significant indirect SNPs localized on GGA4 and GAA1, whereas the opposite pattern was observed for the most significant direct SNPs on GAA3, GGA7, and GGA14, which showed large direct and small indirect effects. Although the overall effects of these SNPs from SEM-GWAS and MTM-GWAS were similar, the use of decomposition allowed us to determine that the trait of interest is affected in different manners. For instance, a given SNP effect may largely act directly on HHP without any mediation by BM and BW, whereas another SNP may be transmitting a large effect through a causal path mediated by BM and BW. Collectively, new insight regarding the direction of SNP effects can be obtained using the SEM-GWAS methodology.

**Table 3.** Six most significant SNPs selected according –log_10_ (p-values) and their effects, using the full recursive SEM (SEM-A75) and MTM (MTMA75). 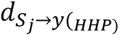, 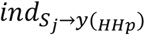, 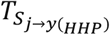 and 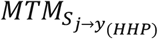, represents, direct, indirect and overall from SEM and MTM effects of j-*th* SNP on HHP.

| | CHR | SNP name | candidate genes | $-\log_{10}(\text{p-values})$ | | | | Type of SNP effect | | | |
| --- | --- | --- | --- | --- | --- | --- | --- | --- | --- | --- | --- |
| | | | | $d_{S_j \rightarrow y(HHP)}$ | $ind_{S_j \rightarrow y(HHP)}$ | $T_{S_j \rightarrow y(HHP)}$ | $MTM_{S_j \rightarrow y(HHP)}$ | $d_{S_j \rightarrow y(HHP)}$ | $ind_{S_j \rightarrow y(HHP)}$ | $T_{S_j \rightarrow y(HHP)}$ | $MTM_{S_j \rightarrow y(HHP)}$ |
| Top SNPs for direct effects | 14 | Gga_rs313620413 | GRIN2A | **7.4242** | 0.1499 | 9.6599 | 7.4525 | -5.7827 | -0.0498 | -5.8326 | -5.78511 |
|  | 7 | Gga_rs16591372 | OLA1 | **7.0868** | 0.2220 | 9.0119 | 6.9783 | -22.5681 | 0.2983 | -22.2698 | -22.3520 |
|  | 3 | Gga_rs15390496 | EPHA7 | **7.0209** | 0.2214 | 8.6122 | 7.0297 | -22.4233 | -0.2149 | -22.6382 | -22.4098 |
|  | 1 | Gga_rs314001234 | ---- | **7.0147** | 1.1067 | 9.0710 | 7.1653 | -26.6538 | -0.9018 | -27.5556 | -26.9360 |
|  | 7 | Gga_rs315626061 | ---- | **6.8300** | 0.3360 | 8.9974 | 6.9529 | 5.1767 | 0.0910 | 5.26783 | 5.22295 |
|  | 7 | Gga_rs316509306 | ---- | **6.8241** | 0.3442 | 8.9952 | 6.9485 | 5.1742 | 0.0928 | 5.267116 | 5.22105 |
| Top SNPs for indirect effects | 4 | Gga_rs316082590 | LOC422264 | 0.7137 | **3.6868** | 0.4754 | 0.5696 | -1.2913 | 0.4505 | -0.84073 | -1.07339 |
|  | 4 | Gga_rs313358833 | LOC422265 | 0.6449 | **3.2345** | 0.4310 | 0.5202 | -1.2067 | 0.4235 | -0.78322 | -1.01618 |
|  | 4 | Gga_rs314615897 | MAEA | 0.1170 | **2.9505** | 0.0474 | 0.0387 | -0.2799 | 0.3853 | 0.105456 | -0.09807 |
|  | 1 | Gga_rs15301842 | ---- | 0.0393 | **2.9408** | 0.1436 | 0.0149 | -0.1301 | 0.5053 | 0.375199 | 0.050463 |
|  | 1 | Gga_rs314551852 | ---- | 0.0632 | **2.8858** | 0.1100 | 0.0065 | -0.2038 | 0.4994 | 0.295514 | -0.02218 |
|  | 1 | Gga_rs317379325 | ---- | 0.1599 | **2.8473** | 0.0070 | 0.0931 | -0.4789 | 0.5000 | 0.021148 | -0.29321 |
| Overall effects | 14 | Gga_rs313620413 | GRIN2A | 7.4242 | 0.1499 | **9.6599** | 7.4525 | -5.7827 | -0.0498 | -5.83262 | -5.7851 |
|  | 1 | Gga_rs314001234 | ---- | 7.0147 | 1.1067 | **9.0710** | 7.1653 | -26.653 | -0.9018 | -27.5556 | -26.9360 |
|  | 7 | Gga_rs315626061 | ---- | 7.0868 | 0.2220 | **9.0119** | 6.9783 | -22.5681 | 0.2983 | -22.2698 | -22.3520 |
|  | 7 | Gga_rs315626061 | ---- | 6.8300 | 0.3360 | **8.9974** | 6.9529 | 5.1767 | 0.0910 | 5.26783 | 5.2229 |
|  | 7 | Gga_rs316509306 | ---- | 6.8241 | 0.3442 | **8.9952** | 6.9485 | 5.1742 | 0.0928 | 5.267116 | 5.2210 |
|  | 7 | Gga_rs15850017 | ZNF385B | 6.6582 | 0.0499 | **8.6397** | 6.6176 | -20.8591 | -0.0718 | -20.9310 | -20.7681 |
| MTM | 14 | Gga_rs313620413 | GRIN2A | 7.4242 | 0.1499 | 9.6599 | **7.4525** | -5.7827 | -0.0498 | -5.8326 | -5.7851 |
|  | 1 | Gga_rs314001234 | ---- | 7.0147 | 1.1067 | 9.0710 | **7.1653** | -26.6538 | -0.9018 | -27.5556 | -26.936 |
|  | 3 | Gga_rs15390496 | EPHA7 | 7.0209 | 0.2214 | 8.6122 | **7.0297** | -22.4233 | -0.2149 | -22.6382 | -22.4098 |
|  | 7 | Gga_rs16591372 | OLA1 | 7.0868 | 0.2220 | 9.0119 | **6.9783** | -22.5681 | 0.2983 | -22.2698 | -22.352 |
|  | 7 | Gga_rs315626061 | ---- | 6.8300 | 0.3360 | 8.9974 | **6.9529** | 5.1767 | 0.0910 | 5.26780 | 5.2229 |
|  | 7 | Gga_rs316509306 | ---- | 6.8241 | 0.3442 | 8.9952 | **6.9485** | 5.1742 | 0.0928 | 5.2671 | 5.2210 |

The bold values are $-\log_{10}(\text{corrected p-value})$ for each type of significant SNP effects categories.

## Discussion

It is becoming increasingly common to analyze a set of traits simultaneously in GWAS by leveraging genetic correlations between traits [37, 38]. In the present study, we illustrated the potential utility of a SEM-based GWAS approach for causal inference and mediation analysis of SNP effects, which has the potential advantage of embedding a pre-inferred causal structure across phenotypic traits [32]. SEM-GWAS, as an extension of standard MTM, accounts for recursive linking of mediating variables that could be either dependent or independent with restriction on a residual covariance. This is a useful approach when multiple mediators influence the final outcomes via either common or distinct biological pathways [39, 40]. SEM-GWAS is achieved by first inferring the structure of networks between phenotypic traits. For this purpose, we used a modified version of the IC algorithm described by Pearl [29] and modified for implementing in quantitative genetics by Valente, et al. [32]. The IC algorithm was used to explore putative causal links among phenotypes obtained from a residual covariance matrix, in a model that accounted for systematic and genetic confounding factors such as polygenic additive effects. It then produced a posterior distribution of partial residual correlations between any possible pairs of variables. Three different causal path diagrams were inferred from HPD intervals of 0.75, 0.85, and 0.95. We observed that the number of identified paths decreased with an increase in the HPD interval value. Only a path connecting BM and BW was present in all HPD intervals considered. Moreover, we found that the partial residual correlation between BM and HHP was weaker than that between BM and BW. This may explain why the path between BM and HHP was not detected with HPD intervals larger than 0.75.

Estimated path coefficients reflect the strength of each causal link, quantifying the proportion of direct and indirect effects of a given SNP or genes on the outcome of interest via the mediator phenotypic traits or the predefined causal pathway between a set of mediators and the target outcome. For instance, a positive path coefficient from BM to BW suggests that a unit increase in BM directly results in an increase in BW. Our results showed that MTM-GWAS and SEM-GWAS were similar in terms of the goodness of fit as per the AIC and BIC criteria. This finding is in agreement with theoretical work of Gianola and Sorensen [11] and Varona, et al. [12] showing the equivalence between models. Thus, MTM-GWAS and SEM-GWAS produced the same marginal phenotypic distributions and goodness of fit values. A similar approach has been proposed by Li, et al. [16], Mi, et al. [41], and Wang and van Eeuwijk [42]. The main difference between our approach and theirs is that they used SEM in the context of standard QTL mapping, whereas our SEM-GWAS is developed for GWAS based on a linear mixed model.

The advantage of SEM-GWAS over MTM-GWAS is that the former decomposes SNP effects by tracing inferred causal networks. Our results showed that by partitioning SNP effects into direct, indirect, and total components, an alternative perspective of SNP effects can be obtained. As shown in Table 3 and Figure 4, direct and indirect effects may differ in magnitude and sign, acting in the same direction or in an antagonistic manner. Note that the total SNP effects inferred from SEMGWAS were the same as the estimated SNP effects from MT-GWAS (Table 3). However, knowledge derived from the decomposition of SNP effects may be critical for animal and plant breeders in breaking unfavorable indirect QTL effects or obtaining better SNP effect estimates than those from MTM-GWAS [e.g., 41].

## Conclusion

SEM offers insights into how phenotypic traits relate to each other. We illustrated potential advantages of SEM-GWAS relative to the commonly used standard MTM-GWAS by using three chicken traits as an example. SNP effects pertaining to SEM-GWAS have a different meaning than those in MTM-GWAS. Our results showed that SEM-GWAS enabled the identification of whether a SNP effect is acting directly or indirectly, i.e. mediated, on given trait. In contrast, MTM-GWAS only captures overall genetic effects on traits, which is equivalent to combining direct and indirect SNP effects from SEM-GWAS together. Thus, SEM-GWAS offers more information and provides an alternative view of putative causal networks, enabling a better understanding of the genetic quiddity of traits at the genomic level.

## Conflict of Interest

The authors do not have any conflict of interest.

## Author’s contributions

MM carried out the study and wrote the first draft of the manuscript. GJMR and DG designed the experiment, supervised the study and critically contributed to the final version of manuscript. GM contributed to the interpretation of results, provided critical insights, and revised the manuscript. BDV and AAM participated in discussion and reviewed the manuscript. MA, AK and RMP contributed materials and revised the manuscript. All authors read and approved the final manuscript.

## Acknowledgment

The first author wishes to acknowledge the Ministry of Science, Research and Technology of Iran for financially supporting his visit to the University of Wisconsin-Madison. Work was partially supported by the Wisconsin Agriculture Experiment Station under hatch grant 142-PRJ63CV to DG.

